# A comparison of the gobiid fauna between a shoal and an island habitat in the central Visayas (Philippines)

**DOI:** 10.1101/006049

**Authors:** Klaus M. Stiefel, Alistair Merrifield, Matt Reed, David B. Joyce

## Abstract

We surveyed the marine gobies of Malapascua island (Philippines), the surrounding islets and the nearby Monad shoal. We found 59 species in 19 genera, including 2 undescribed species of the genus *Trimma*, and 3 geographic and 6 depth range expansions. Furthermore we describe a new type of mimicry between the goby *Koumasetta hectori* and the cardinalfish *Apogon nigrofasciatus*. The comparison of the island versus shoal goby fauna showed a lesser species richness of shrimp-associated gobies at the shoal. This likely reflects the fact that hydrodynamic features of the environment play a dominant role in selecting which gobiid species, or their symbiotic shrimp, will be found in a certain location. We also observed a bias towards hovering species (of the genus *Trimma*) and away from shrimp-associated gobies at greater depths. These findings are in accord with the suspected shift of gobies towards planctotrophy with increasing depth.

We furthermore compare this study to previous surveys of goby faunas, and plot the recorded species numbers against the survey areas. This species-area plot provides support for the notion of high speciation rates in gobies due to their low mobility.

## Introduction

The gobies (Chordata, Vertebrata, Actinopterygii, Teleostei, Perciformes, Gobiidae) are a very species-rich family of small marine and freshwater fishes. The peak of goby (and general marine) biodiversity is found in the “coral triangle”, between Eastern Indonesia, Papua New Guinea and the Philippines [1, 2]. We surveyed the goby fauna in a location within the coral triangle, Malapascua island (11°19’58.35"N, 124° 6’57.40"E), in the central Visayas, Republic of the Philippines. We included in our study both the coastline of Malapascua island proper, a number of surrounding islets within 15 km of Malapascua and the Monad shoal (Fig. 1). The marine area sampled covers approximately 250 km^2^.

**Figure 1:**
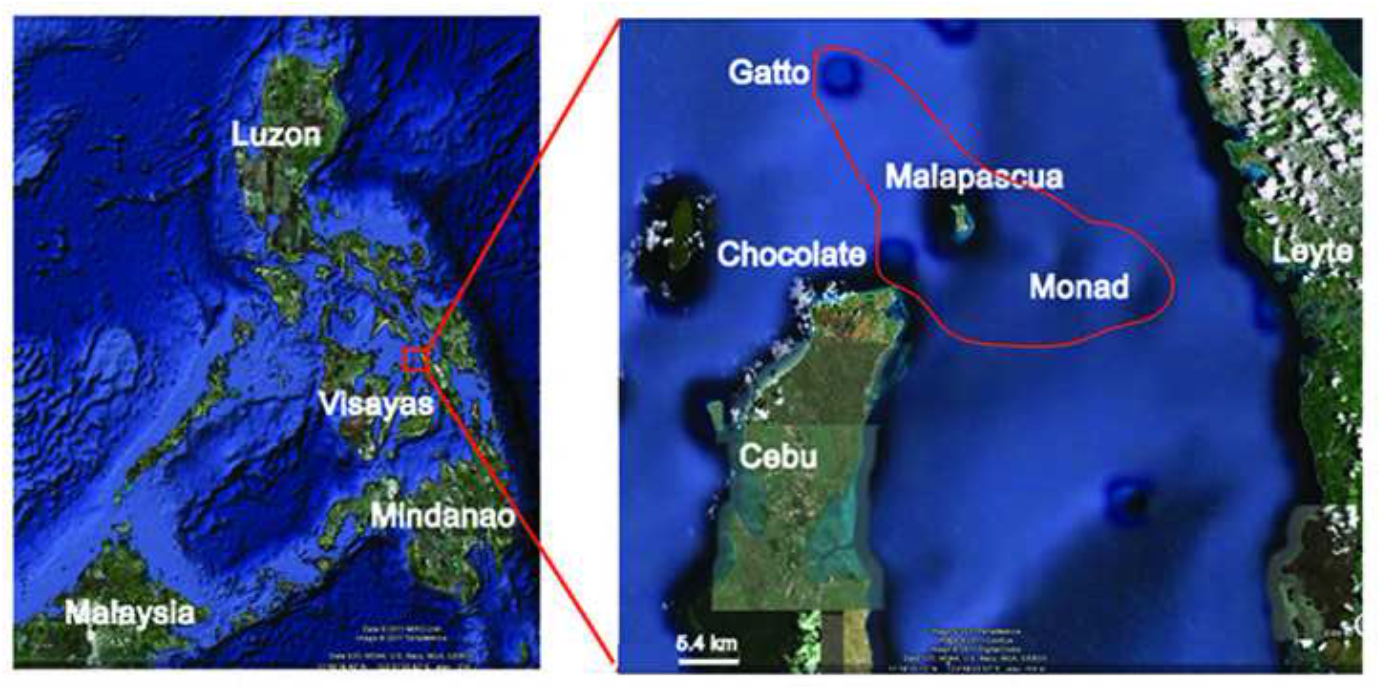
Location of the surveyed sites. Left map: The Philippine archipelago. The area marked in the red box is enlarged in the right map. The region encircled in red is the surveyed area. Maps from Google Earth.

Malapascua island lies 10 km north of the island of Cebu. It has a land area of ∼2.5 km^2^ and is 10 m high at its highest point. It is surrounded by a number of uninhabited islets (Chocolate island, Gatto island) which were also explored in this survey. Marine habitats present include sandy bottoms, seagrass meadows, mangroves and coral reefs. The coral reefs consist of a diverse assembly of hard- and very prolific soft corals and feature diverse invertebrate and fish populations. The reefs transition into sandy planes at a depth of about 55 m. Through coastal development, the mangroves have been reduced to less than 100 m of coastline. There are no flowing streams or freshwater bodies on the island, and only a small area (less than 100 m of coastline) with rocky tidepools.

The nearby Monad shoal rises up from a depth of 93 m to a plateau with an average depth of 20 m, a minimum depth of 13 m, and a maximum depth of 22 m. The surface area of the plateau is ∼2 km^2^. Large parts of it are covered in *Anacropora* and *Xenia* corals, with sandy, rocky and mixed coral areas present as well. Malapascua is home to dive tourism, fueled by frequent sightings of thresher sharks (*Alopias pelagicus*) at the Monad shoal. This dive tourism has led to marine protection efforts including a fairly well enforced fishing ban on Monad shoal, and the coral reef, especially below 25 m, appears very healthy, featuring a dense coral cover and large sea-fans.

The population of Malapascua is ∼ 2000. Anthropogenic influences on the marine fauna in Malapascua include conventional net-based fishing, and unfortunately until recently, fishing using explosives and cyanide (for the aquarium fish market; both observed by the authors). While the dive tourism motivates conservation efforts, it also causes a certain amount of mechanical damage by divers to the reef structure. Even though the effects of destructive fishing on the coral cover are visible at some sites, fish abundance and diversity are still high at these locations. These anthropogenic influences do not necessarily directly influence the goby populations, but will likely have indirect influences via the presence of host corals and predators: A change in the marine food web and invertebrate cover will likely have an influence on the region’s gobiidae.

The intention of this study is fourfold: (1) to conduct a general survey of the gobiid species in the Malapascua area; (2) to compare the gobiid species distributions between two types of oceanic habitats, an island (Malapascua) and a shoal (Monad); and (3) to investigate the depth profile of the gobiid distributions. After reporting on these questions, (4) we compare the results of our study with three other studies of goby faunae, and analyze the species number versus area relationships, a measure with relevance to animal mobility and speciation.

## Materials and Methods

It was our goal to determine the presence of gobiid species, and the depth range of these species, in Malapascua proper and at the Monad shoal. These goals motivated our qualitative sampling approach, as detailed below.

Fish were visually observed during close inspection of the reef during conventional and mixed gas (O_2_/He/N_2_, “trimix”) scuba dives at depths of 1 m to 60 m. Observed species were recorded together with depth, and fish were photographed in most cases. Mangroves, seagrass beds and tidal flats were explored on foot and on snorkel.

We opted for an observational/photographic approach for several reasons: (1) the bottom time during deep dives is limited, and specimen capture takes considerably more time than observation and photography; (2) while environmental damage caused by the use of fish anesthetic is in fact limited, we wanted to set a good example for the still fledging marine conservation efforts in the Philippines by sampling non-invasively.

We do not believe that it is possible to determine the exact numbers or densities of small, sometimes cryptic or semi-cryptic and easily spooked fish, such as gobies, in a non-invasive study like ours. Furthermore, many goby species depend very strictly on the presence of a certain micro-habitat, like a host coral[3]. For instance, the whip coral goby (*Bryaninops yongei*) only occurs on whip corals of the genus *Cirripathes*, and the number of individual fish will strongly depend on the number of corals. Thus, stating a number of individuals per m^2^ only makes limited sense in the case of these highly micro-habitat dependent animals. Therefore, and because we were primarily interested in recording the presence of fish species, not the number of individuals, we decided against transects or area-based fish counting methods. Only occasionally do we state semi-quantitative observations in this study.

It is very likely that we have missed some rare and cryptic gobiid species in our survey. We do however believe that we have observed most non-cryptic gobies in the Malapascua area. Most (57/59) species reported here were sighted more than once, many of them multiple times, indicating a relatively complete sampling of all but the rarest and cryptic species. We hence believe that the list of goby species presented here is of value, and sufficiently complete to support our conclusions.

Fishbase [4], and references [5, 6] were used as the primary references for species identification and expected geographical and depth ranges. For individual species we consulted the specialized literature. The survey was conducted from January 7^th^ to February 6^th^, 2011.

## Ethics Statement

No specimen were collected, hence no sampling permission was necessary.

## Results

### General survey of the Gobiidae

We found 59 species of gobies in 19 genera (Tab. 1). Of these species, 18 were shrimp-associated (as described in [7], Fig. 2), 12 were found in the sand/rubble without symbiotic shrimp (Fig. 3), 2 were dwelling between coral fingers (*Acropora*, Fig. 6), 9 were found under ledges and in small caves (sometimes hovering upside-down, Fig. 4), 9 perching on rocks and encrusting hard corals (Fig. 5), 2 on sponges, and 4 species were found exclusively on soft corals (*Dendronephtia* and *Cirripathes,* Fig. 6). *Koumansetta hectori* interestingly was found in two settings: one, it was hiding between the spines of sea urchins (*Diadema sp.,* Fig. 6), which likely conferred it protection as well as camouflage (the goby’s stripes visually blending in with the spines). In the second setting these fish were found on sandy patches under rock overhangs together with a species of cardinalfish with a very similar coloration (*Apogon nigrofasciatus*, Fig. 7). If this is a functional mimicry has yet to be determined – we are not aware of reports whether *Apogon nigrofasciatus* is poisonous. We furthermore found 1 species in the intertidal zone and 1 species of amphibious mudskipper among the mangrove roots (Fig. 6). The small number of mangrove associated species is likely due to the small remaining area of mangroves on Malapascua island.

**Table 1:**
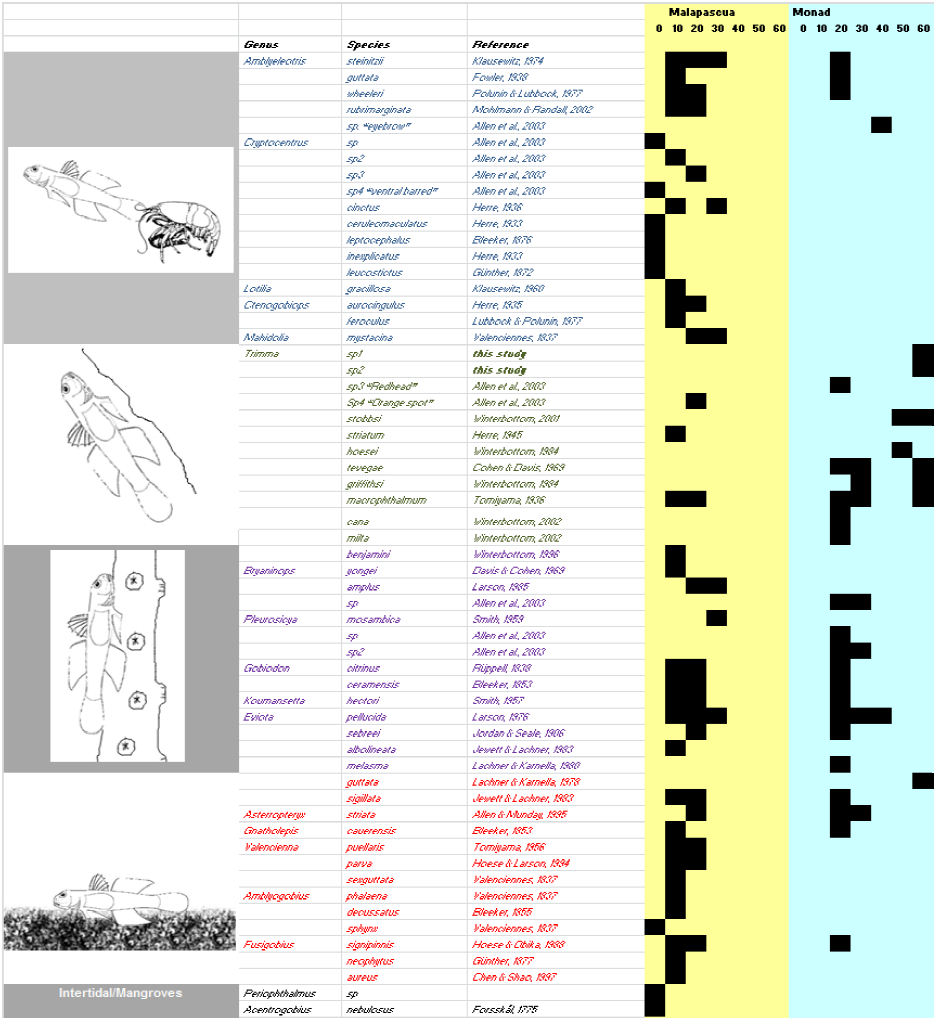
“Gobygram”: Gobies of Malapascua island, surrounding islets and Monad shoal. The black squares in the “Malapascua” (yellow) and “Monad” (blue) columns on the right indicate the observation/photographic record of the respective species at the indicated depth. The species names in citation marks next to “sp” refer to the common names given in [5]. The drawings on the left indicate the ecological specialization of the species. Note the gap in the shrimp associated goby species (empty space in the upper part of the blue section) at Monad shoal.

**Figure 2:**
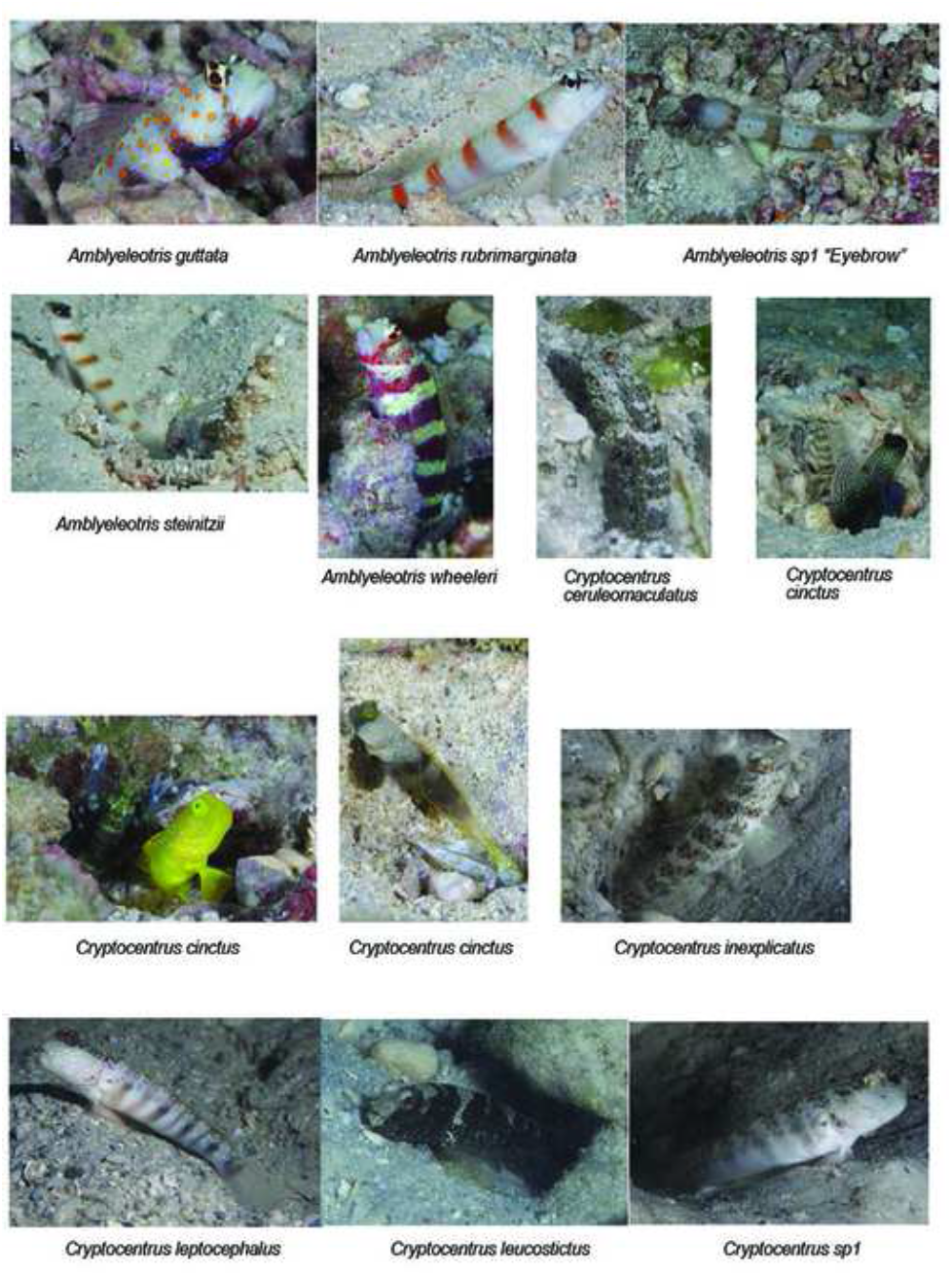
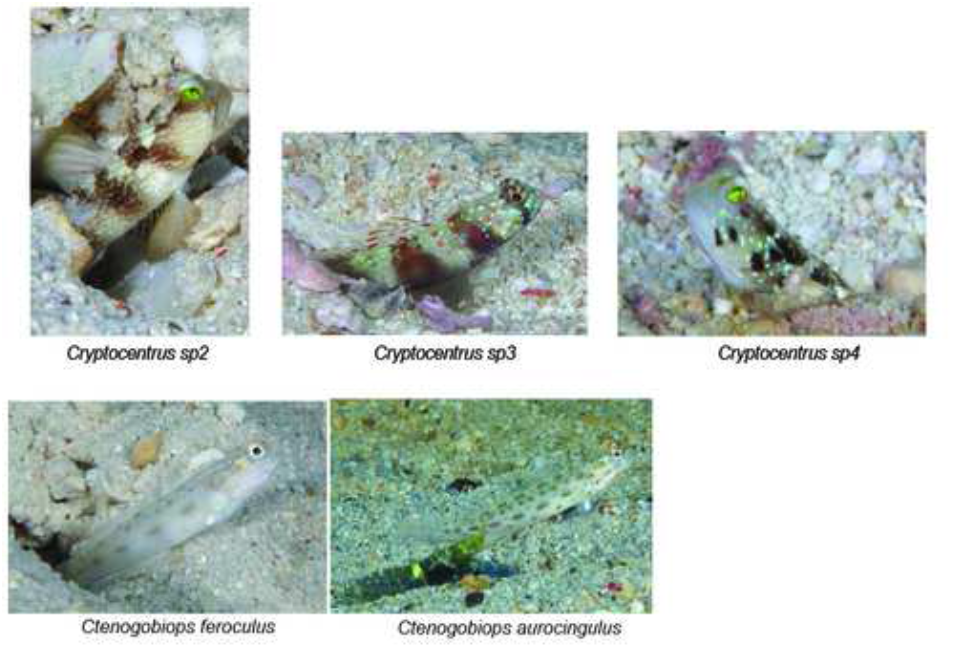
Shrimp-associated gobies. Note the brown, yellow and gray races of *Cryptocentrus cinctus*. Not photographed, but observed: *Lotilia gracilosa*, *Mahidolia mystacina*.

**Figure 3:**
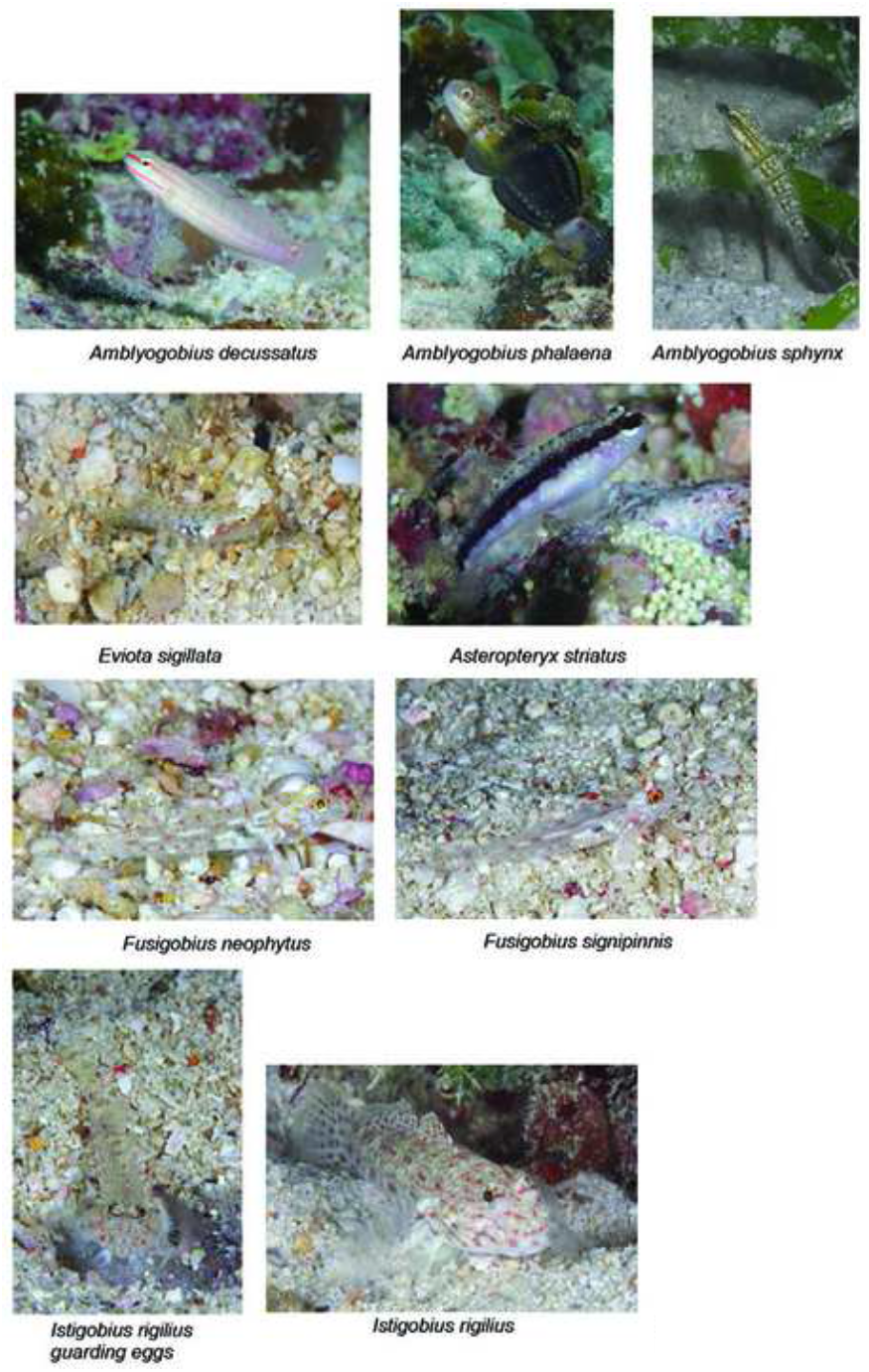
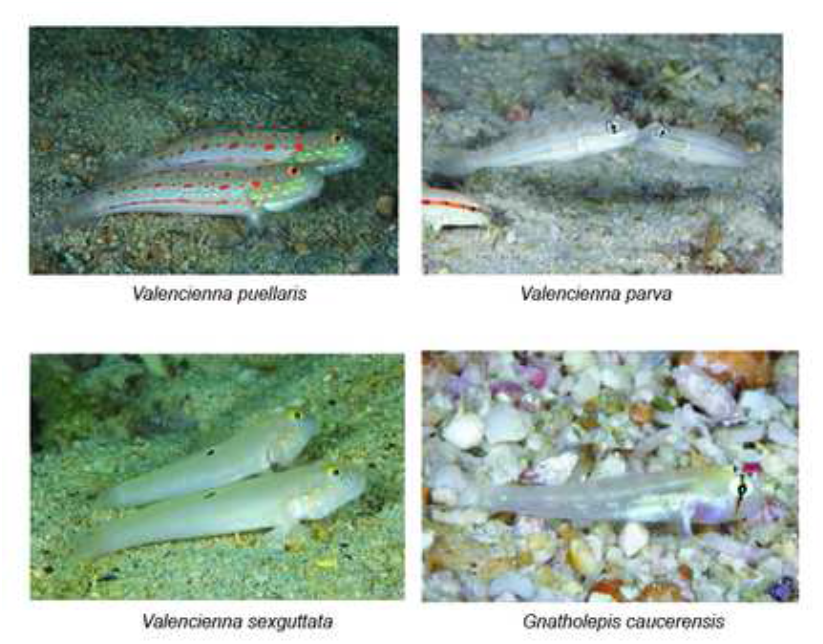
Sand and rubble-dwelling gobies. Note *Istigobius rigilius* guarding its eggs.

**Figure 4:**
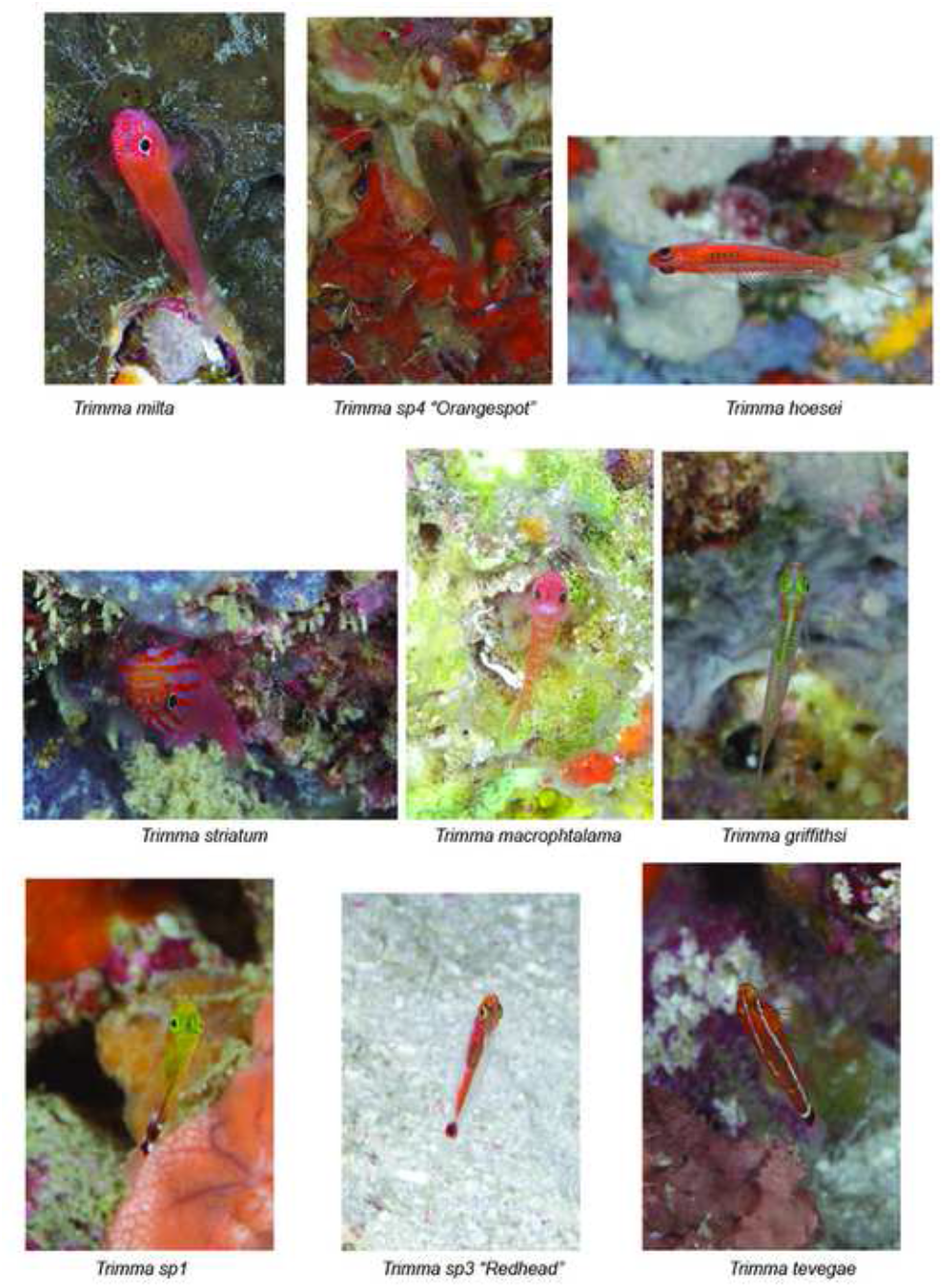
Hovering and cave/ledge dwelling gobies.

**Figure 5:**
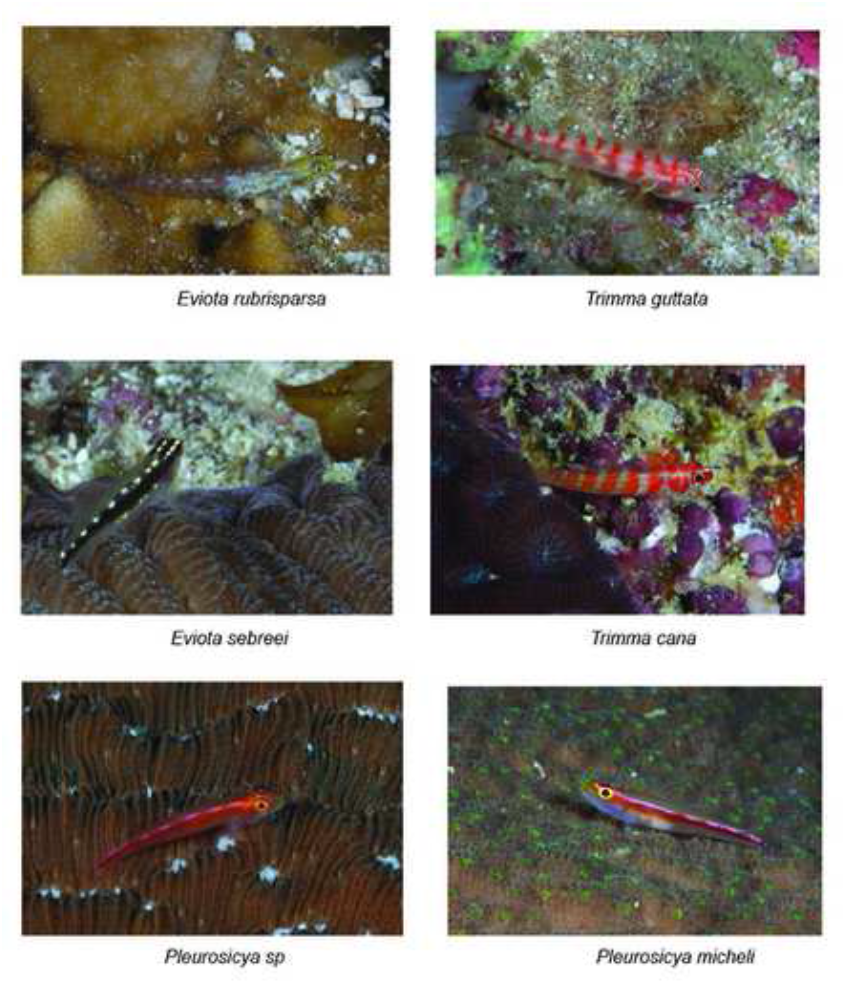
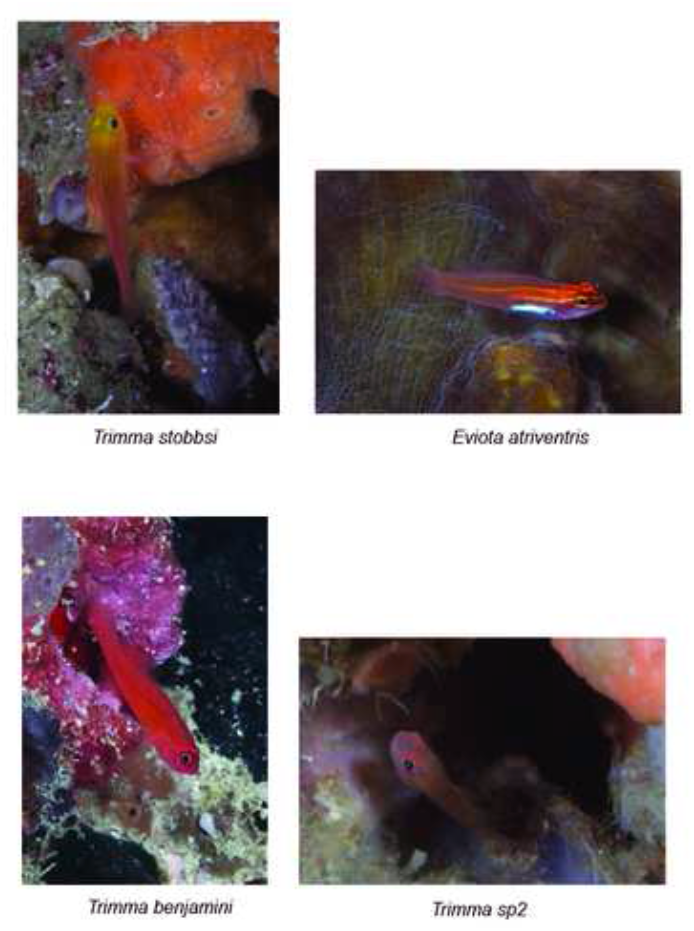
Rock and hard coral perching gobies.

**Figure 6:**
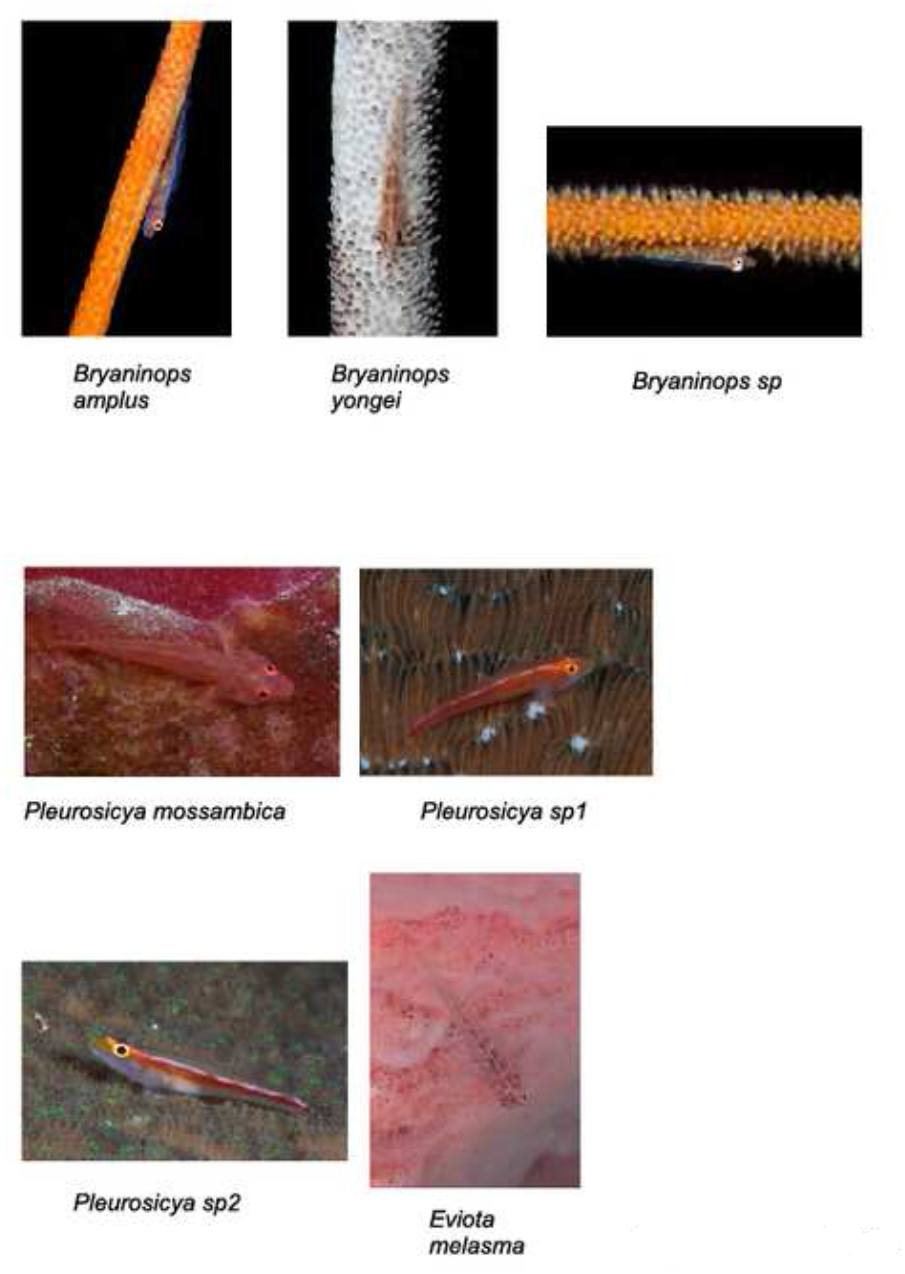
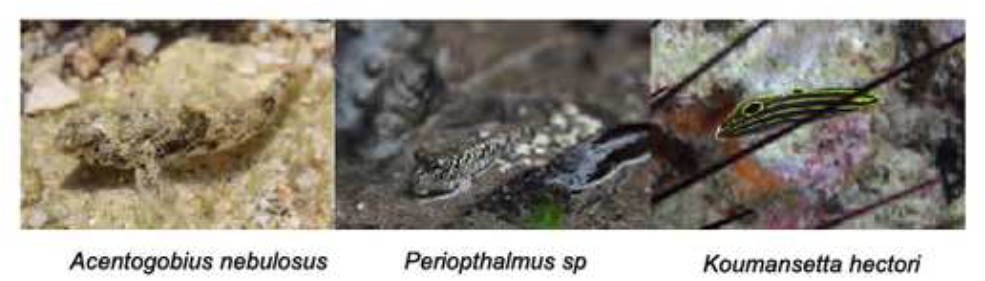
Soft coral, sponge and urchin associated gobies, intertidal goby, mudskipper.

**Figure 7:**
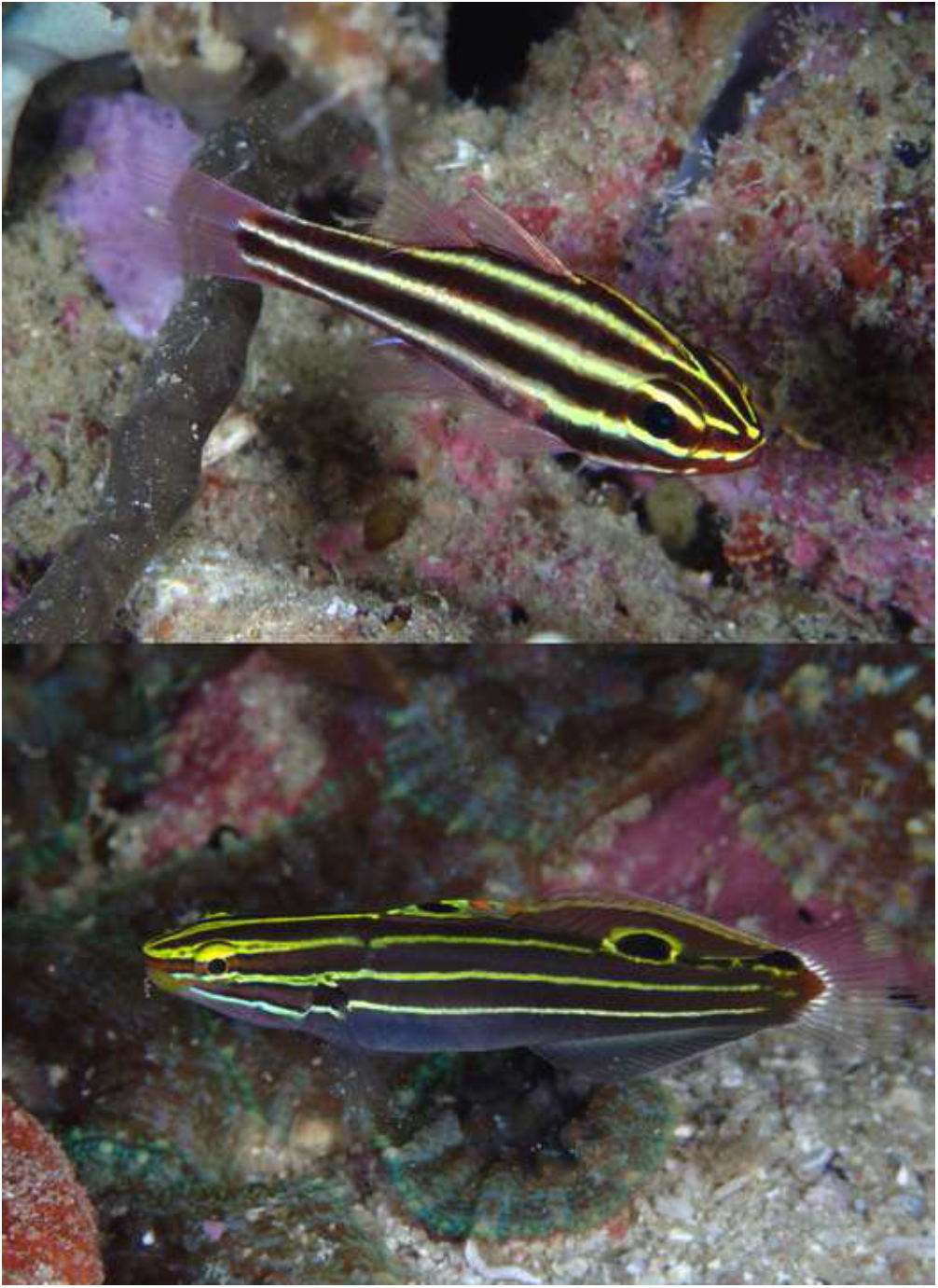
Mimicry between *Koumasetta hectori* and *Apogon nigrofasciatus*. These individuals of the two species were found at a depth of 15 m, in close proximity of each other. These pictures were taken less than 5 m apart within a few minutes. Note the false eye spot on the goby; it appears to face in the opposite direction. Also note the second potential function of the stripes of *K. hectori*, camouflage in-between sea urchin spines (Figure 6–2).

As is well established, most goby species showed a pronounced microhabitat preference[3]. This was less so the case for the most common species found in our study, *Eviota atriventris*. Individuals of this species were found on sand, in coral rubble, on rocks and encrusting corals and in-between *Acropora* coral fingers.

We found geographic range expansions for two species, *Amblyeleotris rubrimarginata*, previously only believed to occur in Indonesia and the GBR[8], and *Amblyeleotris sp*. “eyebrow”, believed to occur only in Indonesia. We also found 6 species significantly deeper than they were believed to occur, namely *Trimma stobbsi* (18-26 m depth range according to fishbase; found in 60 m), *Trimma tevegae* (fishbase: to 40 m; found in 60 m) and *Trimma macrophthalmum* (fishbase: to 30 m; found in 60 m), *Trimma griffithsi* (fishbase: to 32 m; found in 60 m), *Eviota atriventris* (fishbase: to 12 m; found in 40 m) and *Eviota guttata* (fishbase: to 15 m; found in 60 m).

Several species we documented are so far undescribed, but documented in the gray literature. Two photographed species, however, are undescribed and not known from the gray literature either. These are *Trimma* sp1 and *Trimma* sp2 (Ta. 1, Fig. 4), which were both found at 60 m at the Monad shoal. The only records of species similar to *Trimma* sp1 are *Trimma* sp. 34, recorded in the fishpix database and photographed by Y. Sakamoto at 65 m (a similar depth) near Kume-jima in the Ryukyu islands of Japan [9]; a photograph taken in Mactan by Uemura, 15-30 m (R. Winterbottom, personal communication) and records from Vanuatu and Triton Bay, Papua New Guinea (R. Winterbottom, personal communication).

### Island - shoal comparison in regards to gobiid species

We next compared Malapascua proper and its surrounding islets with the Monad shoal (Fig. 8). The main result is a bias towards hovering species, and away from shrimp-associated species at the shoal (gap in the upper right part of Tab. 1).

**Figure 8:**
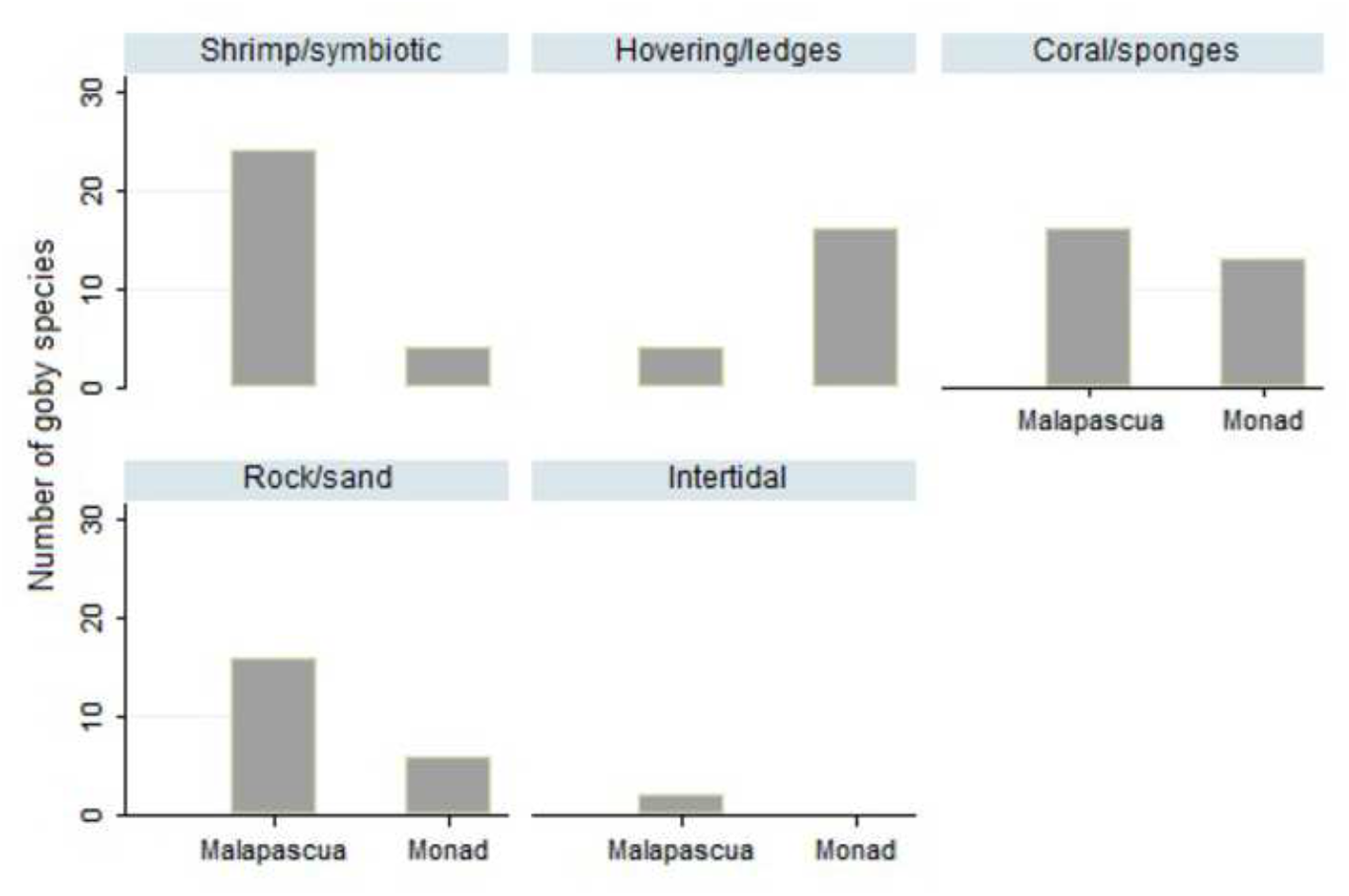
Number of goby species in Malapascua/Monad sorted according to microhabitat.

We found 44 species around Malapascua and associated islets, and 28 species at the Monad shoal. We noted a marked absence of shrimp-associated gobies at the Monad shoal, with a reduced number of species (17 Malapascua / 4 Monad). In contrast, hovering/wall/cave dwelling gobiid species were more prevalent at Monad (4 Malapascua / 8 Monad). Taken together, the relative scarcity of shrimp-associated gobies at the Monad shoal, and the prevalence of hovering *Trimma* lead to a marked difference in genera. In Malapascua, species of *Cryptocentrus* (9 Malapascua / 0 Monad) were more frequent, while in Monad, *Trimma* dominated (4 Malapascua / 9 Monad).

However, only certain depth ranges are present equally at both locations. This ranges spans from 13 m, the top of the Monad shoal, to about 28 m, where the reefs around Malapascua transition into sandy slopes, while the reef wall at Monad continues to drop town near vertically. The very shallow regions, rich in shrimp-associated gobies, are absent at Monad. When we limited our comparison to that depth range, we found a total of 21 species around Malapascua and associated islets, and 23 species at the Monad shoal. Even in this comparison the relative poverty in shrimp-associated gobies at Monad shoal persisted, albeit weaker (7 Malapascua / 4 Monad).

As indicated above, we were restricted to semi-quantitative measurements of population densities in this study. Even so, we estimated the density of shrimp-associated gobies at > 1 m^-2^ around Malapascua island in sand-bottom locations, we typically observed less than 5 individual such gobies during a > 100 m swim-over survey on top of Monad shoal. Assuming a visual inspection by a diver during such a swim of an area ∼1 m to either side, this would amount to an estimated density of < 0.05 m^-2^. As noted, these are semi-quantitative observations, but given the order of magnitude discrepancy we feel that they are worth reporting.

### Influence of depth on gobiid species occurrences

Finally, we compared the gobiid populations at different depths. The main result is a bias towards hovering species at deeper depths.

We found that below 40 m, the vast majority of observed gobies inhabited the rock-ledge, cave and overhang microhabitats (Table 1). Fishes would either perch on the rock or invertebrate cover under these overhangs, often retreating in small burrows when disturbed. Alternatively, they were found in crevices and small caves. Many of these species (belonging to the genus *Trimma.*) hovered in place upside-down. Sandy patches, exposed rocks and encrusting corals and soft corals were curiously devoid of gobies in the 40 – 60 m range. This ecological pattern accounted for the depth-related pattern of gobiid genera. While *Cryptocentrus* is more species-rich in shallower waters, with 9 species in 0 – 20 m and only 1 species in 30 m +, *Trimma* was found to be equally abundant in both zones, with 6 species in 0 – 20 m and 7 species in 30 m +.

### Comparison to previous studies and gobiid mobility

How does the observed number of species, 59, and genera, 19 (in ∼ 250 km^2^) compare with the numbers found in other surveys of gobies in different marine regions? Three published studies in the coral triangle surveyed the gobies of Lizard Island (∼ 40 km^2^, 30 species, [10]), Singapore (∼ 1925 km^2^, 149 species, [11]) and the Papuan Bird’s Head Peninsula (∼ 50 000 km^2^, 308 species, [12, 13]). These studies are significantly different from ours in that they sampled invasively and/or in different depth ranges (Tab. 2). Nevertheless, we believe that the errors in species numbers will not of orders of magnitude, which is sufficient for the comparison attempted below. The slope of a log/log plot of species number against the area these species occur is an indication of their mobility and dispersal [14]. Larger slopes indicate lower mobility and dispersal, and a greater potential for speciation by geographical isolation.

**Table 2:**
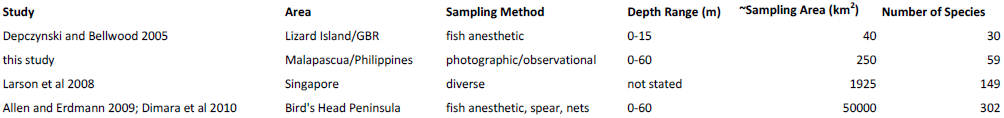
Sampling regions, methods, depth ranges, sampling areas and number of species found in studies from the literature and in this study.

Across the studies, the number of goby species monotonically increases with increasing sampled area, with a slope of 0.327 in a species/area log/log plot (Fig. 9). This value is close to the value expected for terrestrial species on islands [14]. This indicates a weak connectedness between goby populations in the Indo-Pacific. This high *z*-value is consistent with a high territoriality and low mobility of these small, bottom-dwelling, demersally spawning fishes across the expanses of open ocean segregating the reefs of the Indo-Pacific. This low connectedness is also in good accord with the intensive speciation [15] and the resultant high number of Indo-Pacific gobiid species.

**Figure 9:**
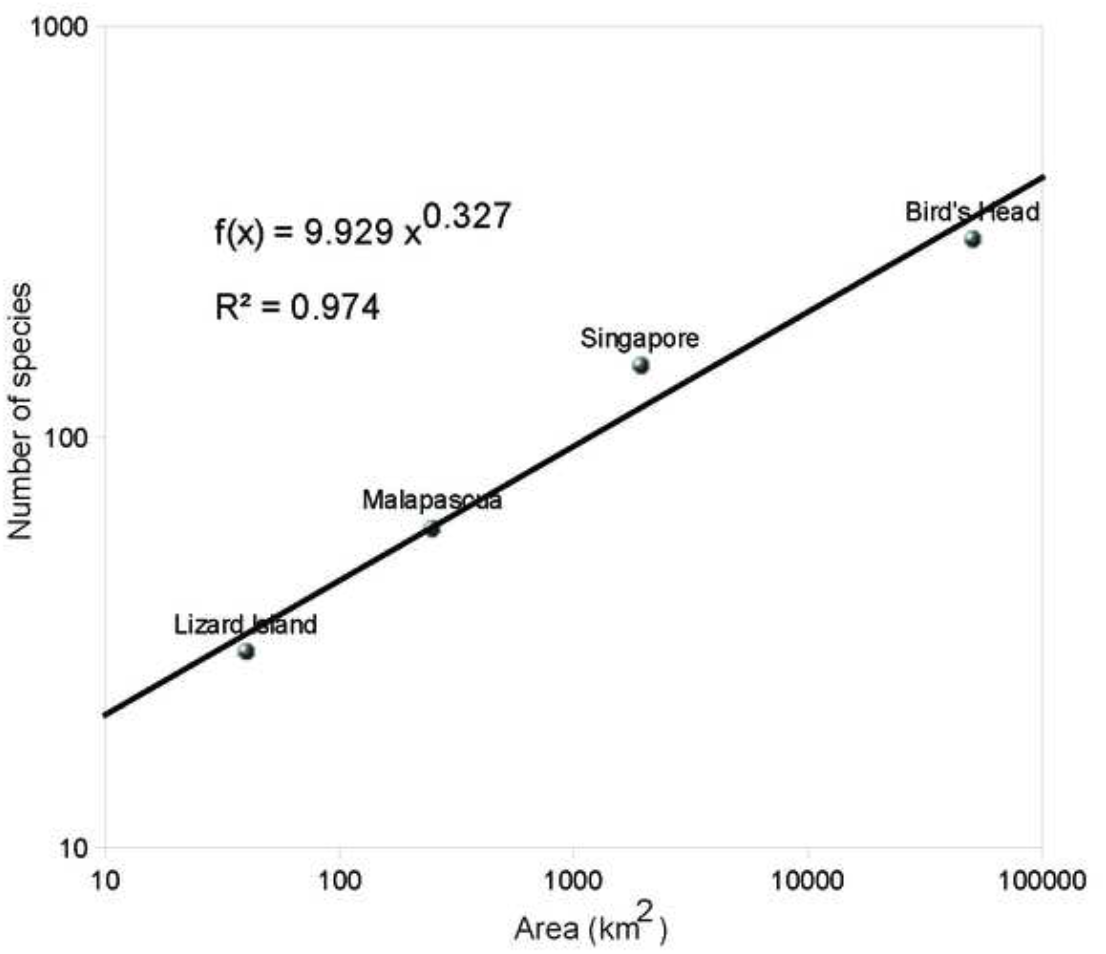
Species – area curve for marine gobies. Plotted are the number of species against the estimated survey area from this study (∼ 250 km^2^, 59 species), a study of the gobies of Lizard Island (∼ 40 km^2^, 30 species), of Singapore (∼ 1925 km^2^, 149 species) and the Papuan Bird’s Head Peninsula (∼ 50 000 km^2^, 302 species). The equation refers to the slope and quality of fit of the linear regression line of this log-log plot. The slope, or *z*-value, is an indicator of the connectedness of populations, with higher *z*-values indicating a lower connectedness.

## Discussion

### Patterns of gobiid distribution

In our study, we observed two noteworthy patterns of gobiid species distributions: More shrimp-associated gobies near the islands, and more hovering gobies at a shoal and at depth. How can these patterns be explained?

Our comparison of the gobies of the island of Malapascua and nearby smaller islands with the Monad shoal, at comparable depth ranges, showed a marked absence of shrimp-associated gobies, despite an ample presence of the sand microhabitats these fishes prefer. Likely, the explanation for this observation lies in the predominance of hydrodynamical features over microhabitat availability in determining the occurrence of gobiid species [10]. The top of the Monad shoal (at 13 - 22 m) is a habitat with rather strong currents - this is also apparent from the presence of a number of current-adapted reef fishes, such as *Pterocaesio* sp.*, Caesio* sp. and *Genicanthus lamarck.* Thus, most likely, the strong currents on top of the Monad shoal prevents shrimp-associated gobies from establishing themselves. Most likely, the strong currents will reduce the stability of the burrows dug by the symbiotic shrimps. Alternatively, the absence of shrimp-associated gobies could be due to an absence of Alpheid symbiotic shrimps [16]. The effect of the physical conditions could then be indirect, via an effect on the shrimps.

When sampling deeper zones, below 30 m, we found an increased number of gobies hovering under ledges and perching on the steep reef walls (mostly *Trimma sp.*). In contrast, the number of gobies living in sandy areas and on soft corals significantly decreased. What is the reason for the change in micro-habitat usage of gobies at deeper depths? It is known that detritus, a main food source for sand-living gobies, decreases as a function of depth [17]. At deeper depths, planctivoric gobies are thought to prevail (D. Bellwood, personal communication). The presence of hovering and reef-wall perching, presumably planctivorous, *Trimma* also point to such trends.

## Author Contributions

K.M.S. conceived the study, sampled and analyzed the data, and wrote the paper. M.R and D.B.J. sampled the data, A.M. analyzed the data.

## Acknowledgements

We thank Jim Scarbourough and Drs. Robert Warner, Satoshi Mitarai, Richard Winterbottom and David Bellwood for helpful discussion and pointers to the relevant literature. We especially thank Dr. Winterbottom for help with the species identification, and the reviewers of a previous version for corrections.

